# Seeding paradise: germination ecophysiology of *Xyris paradisiaca* Wand. (Xyridaceae), an endangered endemic species from Central Brazil

**DOI:** 10.64898/2026.09.24.754203

**Authors:** Carlos A. Ordóñez-Parra, Mateus R. B. Bispo, Lara Amaral-Garcia, Rafael Figueiredo, Tales E. B. Almeida, Claudomiro A. Cortês, Queila S. Garcia

## Abstract

Seed ecophysiology is essential for understanding plant regeneration and developing effective conservation and ecological restoration strategies, yet knowledge remains scarce for most threatened Cerrado species. We characterised the seed germination niche, desiccation tolerance and storage behaviour of *Xyris paradisiaca* (Xyridaceae), an Endangered species endemic to Central Brazil, to assess whether its restricted distribution is associated with narrow germination requirements and to inform seed-based conservation. Germination was tested across light regimes, constant temperatures, decreasing water potentials and short-duration heat shocks, while thermal- and hydro-time models were used to quantify thermal and hydric thresholds. Seeds exhibited an absolute light requirement but broad abiotic tolerances. Germination remained >87% between 15 and 40 °C, with estimated cardinal temperatures of T_b_ = 8.94 °C, T_o_ = 33.13 °C and T_c_ = 45.53 °C. Germination remained similar to the control down to −0.6 MPa and exceeded 40% at −1.0 MPa, with a median base water potential of Ψ_b_= –1.03 MPa. Germination also remained high after 1-min heat shocks up to 200 °C. Seeds were desiccation tolerant, with 100% germination after drying and a viability loss index of −0.04, and germination remained >90% after 24 months of ambient storage. Thus, the highly restricted distribution of X. paradisiaca is not driven by a narrow physiological germination niche. Its broad environmental tolerances, desiccation tolerance and favourable storage behaviour also highlight its unexplored potential for seed-based restoration, propagation and ex situ conservation.

## INTRODUCTION

Seed ecophysiology provides a fundamental basis for understanding plant regeneration and informing biodiversity conservation and management. By characterising how seeds respond to environmental cues, seed ecophysiological studies help define the germination niche of a species (i.e., the range of conditions under which germination can occur) and identify potential environmental filters acting during this critical stage of recruitment (Grubb, 1977; Donohue *et al*., 2010; Dalziell *et al*., 2022). Moreover, such information can guide ex situ seed conservation and seedling production for conservation and restoration purposes, as well as the design of field sowing and translocation strategies (Larson *et al*., 2023; Liyanage *et al*., 2023; CruzOTejada *et al*., 2026). Seed ecophysiology therefore provides an important link between understanding the environmental conditions that regulate plant regeneration and developing effective conservation and restoration strategies.

This is particularly relevant in the Brazilian Cerrado, the world’s most species-rich savannah, which harbours more than 13,000 plant species, nearly half of them endemic (The Brazil Flora Group, 2018). Despite this exceptional diversity, approximately half of its original extent has been converted to agriculture and cattle ranching (Bustamante and Reis, 2026). Accordingly, the Cerrado is recognised as a global biodiversity hotspot and one of the world’s most threatened tropical biomes nowadays (Myers *et al*., 2000; Strassburg *et al*., 2017). Despite its ecological importance and increasing conservation and restoration needs, the processes that sustain natural regeneration remain comparatively understudied in the Cerrado, particularly with respect to seed biology and germination ecology, representing a major knowledge gap for the biome (Buisson *et al*., 2019, 2021).

Most of the exceptional diversity of the Cerrado flora is concentrated in the grassy-shrubby ground layer, which hosts nearly three herbaceous and subshrub species for each woody species (Castro *et al*., 1999; Silva *et al*., 2024). Among the most notable families within the herbaceous layer is Xyridaceae, with 131 species being recorded in the Cerrado, 116 (88.5%) of which are endemic to Brazil, and *Xyris* being its most diverse genus (Silva *et al*., 2024). Most Xyridaceae species occur in seasonally or permanently wet grasslands (locally known as *campos úmidos*) and moist rocky habitats associated with *campos rupestres* (Alves *et al*., 2015).

Previous studies in Neotropical Xyridaceae have shown that *Xyris* species produce very small (<2 mm), low mass (∼0.025 mg) and non-dormant seeds that typically exhibit a strong or absolute light requirement for germination, and germinate at temperatures between approximately 15 and 30 °C (Garcia *et al*., 2020). In contrast, responses to declining water potential and seed storage behaviour remain poorly characterised for the genus, while the few studies addressing fire-related cues indicate considerable tolerance to heat and responsiveness to smoke-derived compounds (Oliveira *et al*., 2018; Motta *et al*., 2024). However, most of these studies have been done with species from *campo rupestre* vegetation in south-eastern Brazil. These ecosystems have distinctive environmental and evolutionary histories (Silveira *et al*., 2016), raising the question of whether germination patterns documented in them are representative of *Xyris* species occurring under contrasting environmental conditions elsewhere that provide different germination cues. This is especially relevant for narrowly distributed endemic species, whose restricted geographical ranges may reflect specialised regeneration requirements (Thompson and Ceriani, 2003).

In this study, we provide a comprehensive assessment of the germination ecophysiology of *Xyris paradisiaca* Wand., an Endangered species endemic to Central Brazil. First, we assessed the germination responses of *X. paradisiaca* to four environmental factors relevant to regeneration in the Cerrado: light, temperature, water potential and heat shocks. We also evaluated seed desiccation tolerance and changes in viability during storage. Based on the strong light requirement reported for congeners, we predicted that *X. paradisiaca* seeds would require light for germination. If its restricted geographical distribution is associated with specialised germination requirements, we further predicted narrow thermal and hydric germination niches. Given the fire-prone nature of the Cerrado vegetation and the heat tolerance reported for other *Xyris* species, we expected that seeds could tolerate short-duration heat shocks. Finally, based on their small size and the storage behaviour reported for congeners, we predicted that seeds would tolerate desiccation and retain high germinability during storage.

## MATERIALS AND METHODS

### Study species

*Xyris paradisiaca* Wand. is a perennial, tuft-forming herb endemic to Goiás and the Federal District, Brazil, where it occurs in *campos úmidos* and *campos rupestres* at elevations of 1,000–1,500 m a.s.l. Reproductive individuals produce scapes reaching 80–100 cm in height, each terminating in a broadly ovoid, multiflowered spike bearing yellowish to light-brown bracts and more than 20 yellow flowers (Fig. 1). Flowering occurs mainly from March to August (Wanderley, 1989, 2009).

**Figure 1.**
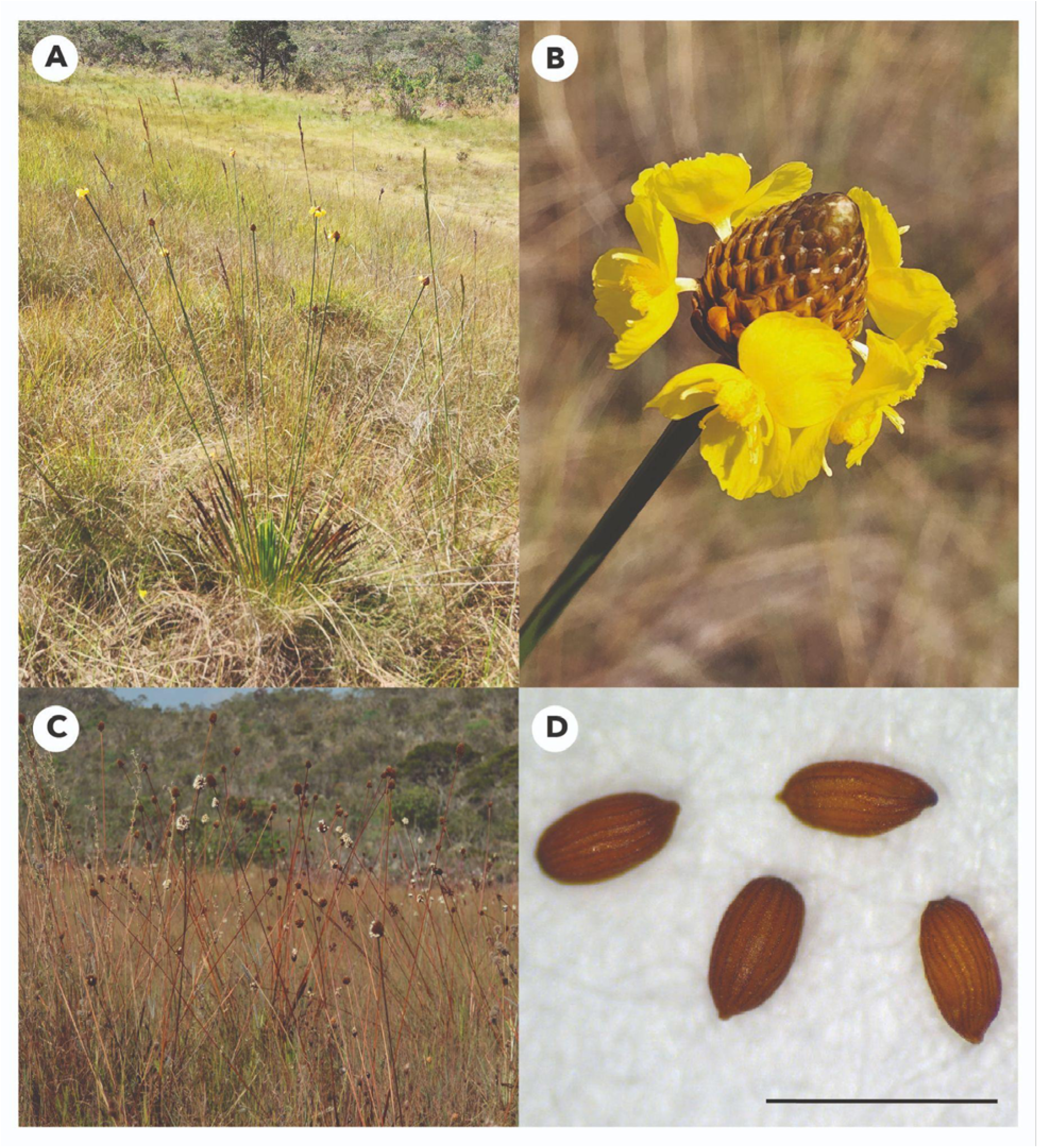
Habit and reproductive structures of *Xyris paradisiaca*. (A) Flowering individual in a *campo úmido*; (B) inflorescence with yellow flowers; (C) individuals bearing mature seeds; and (D) seeds (scale bar = 1 mm). Photos by Carlos A. Ordóñez-Parra (A, B and D) and Lara Amaral-Garcia (C).

*X. paradisiaca* is classified as Endangered in Brazil because it has a very small range of occurrence and area of occupancy and is known from fewer than five threat-defined locations (Araújo *et al*., 2014). Although some populations occur within the Parque Nacional da Chapada dos Veadeiros (PNCV), the habitats of the species are exposed to several increasing pressures. Across the Cerrado, agricultural expansion has been associated with changes in fire frequency and intensity (Abreu *et al*., 2022; Segura-Garcia *et al*., 2025), which may favour the abundance of invasive grasses (Damasceno and Fidelis, 2020). Moreover, unregulated tourism in the region (Araújo *et al*., 2014) and selective harvesting of mature flowering individuals for use in dried “everlasting-flower” arrangements have been reported as additional pressures on reproductive success (Martinelli and Moraes, 2013). Together, these pressures are inferred to contribute to continuing declines in the species’ extent of occurrence, area of occupancy, habitat extent and quality, and the number of mature individuals (Araújo *et al*., 2014).

### Seed collection and germination experiments

In September 2024, mature inflorescences of *X. paradisiaca* were collected from a wet grassland located in the eastern sector of the PNCV. Inflorescences were sampled from 50 spatially separated reproductive individuals and pooled after collection. Seeds were subsequently separated from the surrounding floral structures using a series of sieves with different mesh sizes and stored in sealed plastic tubes enclosed in paper bags, in darkness at room temperature, until use in the experiments. The collection was authorised under SISBIO permit no. 93069, and a voucher specimen was deposited in the Herbarium of the Universidade Federal de Minas Gerais (BHCB, no. 221862). Seeds were stored under ambient laboratory conditions (∼28 °C) for approximately 1, 3, 4 and 12 months before the temperature and light, water potential, heat-shock and desiccation tolerance experiments, respectively.

For each experimental treatment, four replicates of 25 seeds were placed in 60-mm-diameter Petri dishes lined with two layers of filter paper. Unless otherwise stated, seeds were incubated under standard germination conditions of 25 °C, a 12-h photoperiod with a photosynthetic photon flux density of approximately 40 μmol m□² s□¹, and filter paper moistened with distilled water and re-moistened as necessary to prevent the substrate from drying during the experiments. A temperature of 25 °C was selected as the standard incubation temperature because it lies within the range favourable for germination in the *Xyris* species studied to date (Garcia *et al*., 2020). All germination experiments were monitored daily for 30 days. Protrusion of the embryonic axis was used as the germination criterion, and germinated seeds were removed from the Petri dishes. At the end of each experiment, the viability of non-germinated seeds was assessed using a tetrazolium test. Seeds were longitudinally bisected and placed on filter paper moistened with 3 mL of a 1% 2,3,5-triphenyl tetrazolium chloride solution for 48 h at 25 °C in opaque germination boxes. Seeds whose embryos stained uniformly pink or red were classified as viable, whereas seeds with partially stained or nonstained embryos, or with embryos showing grey discolouration, were classified as non-viable.

### Effects of temperature and light

To evaluate the effects of temperature on seed germination, seeds were incubated in germination chambers at constant temperatures of 10–45 °C at 5 °C intervals under the standard 12-h photoperiod described above. We focused on constant temperature regimes because previous studies have shown that daily temperature fluctuations do not enhance germination in *Xyris* species (Abreu and Garcia, 2005), consistent with broader patterns reported for species from Brazilian open ecosystems (Ordóñez-Parra *et al*., 2025, 2026). To evaluate the effects of light on germination, seeds were incubated at 25 °C under either the standard illuminated conditions described above or continuous darkness. The illuminated treatment corresponded to the 25 °C treatment of the temperature experiment, whereas continuous darkness was achieved by enclosing the Petri dishes in black germination boxes. The illuminated treatment was monitored daily, as described above, whereas germination in the dark treatment was scored under a green safe light at 7-day intervals to minimise seed exposure to germination-promoting light (Abreu and Garcia, 2005).

### Effects of water potential

To evaluate the effects of reduced water potential, Petri dishes were moistened with 3 mL of polyethene glycol 6000 (PEG 6000) solutions adjusted to water potentials of −0.2, −0.4, −0.6, −0.8 and −1.0 MPa. PEG 6000 concentrations were calculated for 25 °C according to Michel (1983). These treatments were compared with the distilled-water control corresponding to the 25 °C treatment of the temperature experiment. Petri dishes were sealed with plastic film to reduce evaporative water loss. Every three days, the solutions in all treatments were replaced to maintain the target water potentials.

### Effects of heat shocks

To evaluate the effects of heat shocks, dry seeds from each replicate were placed in a glass Petri dish and exposed to 80, 100, 120, 150 or 200 °C for 1 min in a preheated drying oven. Each replicate was treated separately, with the 1-min exposure beginning when the Petri dish was placed in the oven. The oven was allowed to return to the target temperature before the next replicate was treated. The heat-shock treatments were compared with a nonheated control. The treatments were selected to represent the thermal conditions that seeds may experience during Cerrado fires, from lower temperatures below ground to higher temperatures at or near the soil surface (Zupo *et al*., 2022). After the heat shock treatment, seeds were incubated under the standard germination conditions described above.

### Seed desiccation tolerance assessment

To evaluate whether seeds tolerate desiccation, we applied an adaptation of the 100-seed test of Pritchard *et al*., (2004) as modified by Mattana *et al*., (2020). Briefly, seeds were divided into three sublots: fresh, dried and moist-stored seeds. Two replicates of 13 fresh seeds were germinated without further treatment to determine baseline germination percentage. Seeds assigned to the drying treatment were progressively dried at room temperature for successive 7-d periods using seed-to-silica-gel mass ratios of 1:1 and 1:4, followed by storage over excess silica gel until a seed-equilibrated relative humidity of approximately 15% was recorded, as measured using a thermohygrometer inside the sealed container. Meanwhile, moist-stored seeds were maintained in a semi-closed container at room temperature for the same period. After treatment, dried and moist-stored seeds were allowed to equilibrate under room conditions for 24 h. Two replicates of 13 seeds from each sublot were then incubated under the standard germination conditions described above. Germination percentage of dried seeds was compared with that of both fresh and moist-stored seeds. Fresh seeds provided a baseline estimate, whereas the moist-stored treatment allowed losses caused by desiccation to be distinguished from those associated with the duration and conditions of storage. The remaining seeds were used to estimate seed water content in the fresh seeds (10 seeds) and at the end of the dry and moist storage (6 each) using the ISTA (2007) protocol.

### Effects of storage under ambient laboratory conditions

To evaluate the effects of seed storage on germination and viability, separate seed lots were stored for 3, 6, 9, 12, and 24 months in sealed plastic tubes inside paper bags, kept in darkness at room temperature, and subsequently incubated under the standard germination conditions described above. Germination of freshly collected seeds was used as the 0-month baseline, and the viability of non-germinated seeds at each storage interval was assessed using the tetrazolium test described above. These simple, low-technology storage conditions were selected to represent a readily accessible method that could be used by seed collectors and restoration practitioners in the Cerrado.

### Data analyses

To test the effects of temperature, water potential, heat shocks and storage time on final germination percentage and seed viability, we fitted two separate Generalised Linear Models (GLMs) for each experiment, both with a binomial error distribution and logit link. Treatment was included as a categorical fixed effect in each model. For germination, the response was adjusted for seed viability within each replicate. The total number of viable seeds was calculated as the sum of germinated seeds and non-germinated seeds classified as viable by the tetrazolium test; therefore, models were fitted to the numbers of germinated and viable but non-germinated seeds. For seed viability, models were fitted to the numbers of viable and non-viable seeds in each replicate, with viable seeds comprising both germinated and non-germinated seeds classified as viable by the tetrazolium test.

Since some treatments exhibited complete (100%) or null (0%) germination, which can result in (quasi-)complete separation in binomial GLMs, models were fitted using bias-reduced estimation implemented in the *brglm2* package (Kosmidis, 2026). Model assumptions were evaluated using simulation-based residual diagnostics from the DHARMa package (Hartig, 2026), including assessments of residual uniformity, dispersion, outliers and residual patterns against fitted values and treatment levels. The overall effect of each treatment was evaluated using Wald chi-squared tests as implemented in the car package (Fox and Weisberg, 2019). When a significant overall treatment effect was detected, post hoc pairwise comparisons were conducted using Šidák-adjusted contrasts via the *emmeans* package (Lenth and Piaskowski, 2026).

The germination responses of *X. paradisiaca* to temperature and water potential were characterised using thermal- and hydro-time-to-event models, respectively, as implemented in the *drcSeedGerm* package (Onofri *et al*., 2018; Onofri, 2023). Daily germination counts were expressed as interval-censored time-to-event data, with each germination event assigned to the interval between consecutive observations. Thermal time models were used to estimate the base temperature (T_b_), optimal temperature (T_o_), ceiling temperature (T_c_) and the thermal time constant (θ□). Hydro-time models were used to estimate the median base water potential for germination (Ψ_b_) and the hydro-time constant (θ□). Reported thermal- and hydro-time constants and base water potential corresponded to the 50th germination percentile. For each environmental factor, all candidate thermal- or hydro-time models available in the *drcSeedGerm* package were fitted, and among models that converged, the model with the lowest Akaike Information Criterion (AIC) was selected as the best-supported model (Akaike, 1974).

To characterise the effect of desiccation on germination, we calculated the Viability Loss Index (VLI) as follows:

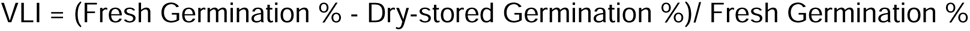

VLI values theoretically range from 0, indicating complete desiccation tolerance, to 1, indicating complete loss of germination following desiccation; although negative values can occur when dried seeds germinate more successfully than fresh seeds. To assign the seed lot to a qualitative category, VLI values were classified as desiccation tolerant (VLI ≤ 0.05), potentially desiccation tolerant (0.05 < VLI ≤ 0.50), potentially desiccation sensitive (0.50 < VLI ≤ 0.95) or desiccation sensitive (VLI > 0.95), based on the thresholds proposed by Mattana et al., (2020). All analyses and visualisations were conducted in R version 4.5.1 (R Core Team, 2025).

## RESULTS

*X. paradisiaca* seeds exhibited an absolute requirement for light, with seeds germinating only under the 12-h photoperiod but not in continuous darkness at 25 °C. Temperature had a significant effect on germination (df = 7, □^2^ = 123.76, p < 0.001). Germination percentage remained high (>87.2%) between 15 and 40 °C, and little or no germination occurred at 10 or 45 °C (Fig. 2A). Temperature also significantly affected seed viability (df = 7, □^2^ = 49.51, p < 0.001), being the highest between 15 and 35 °C (>92.7%) and declining outside this range, reaching 78.8% at 10 °C, 74.4% at 40 °C and 77.0% at 45 °C (Table 1). According to the best-supported thermal-time model, the thermal window for germination extended from T_b_ = 8.94 °C to T_c_ = 45.53 °C, with a T_o_ of 33.13 °C. The estimated thermal-time constant was θ□ = 70.02 °C d.

**Figure 2.**
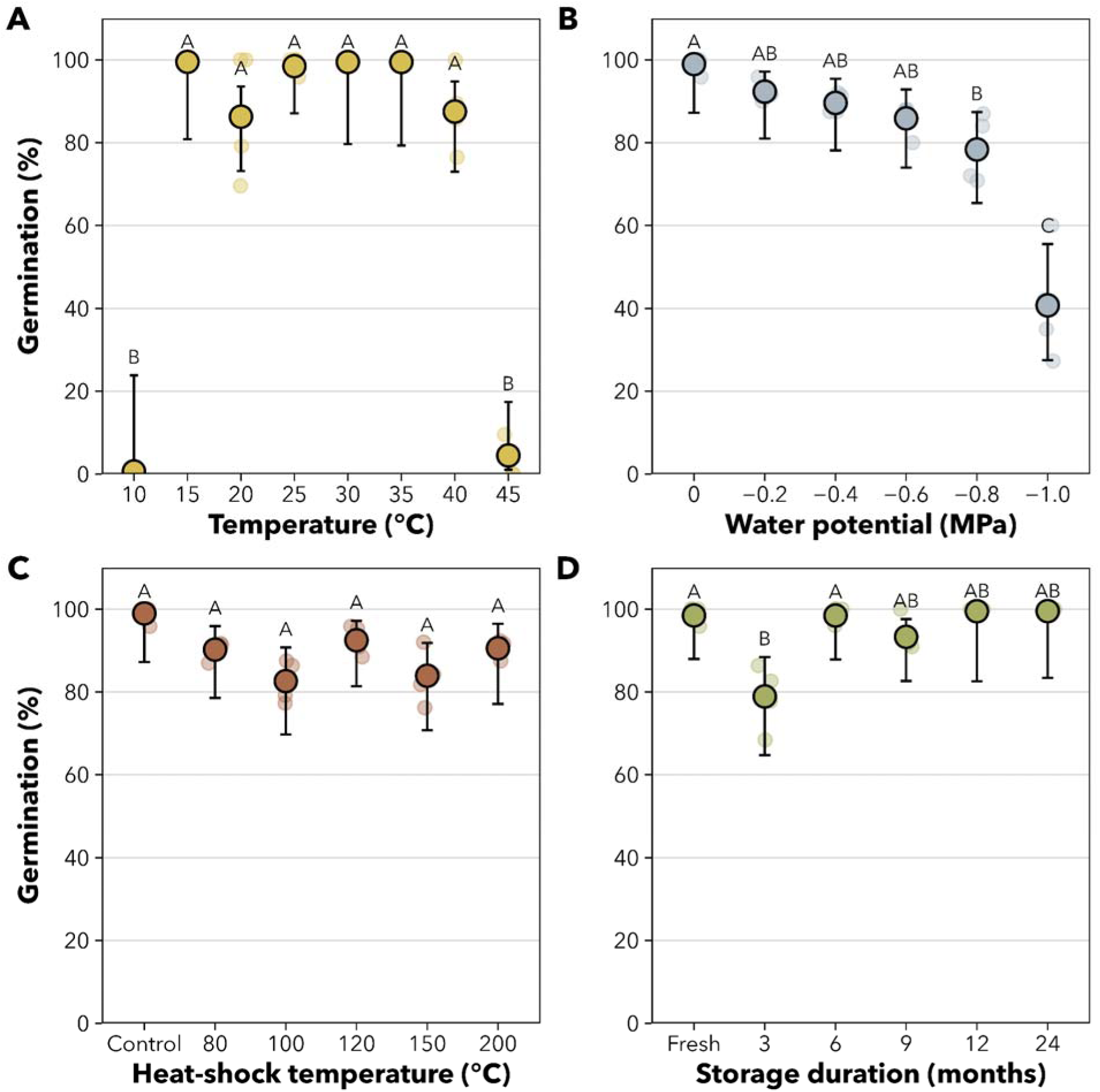
*Xyris paradisiaca* germination responses to environmental conditions and storage. Germination percentage under (A) constant temperatures, (B) decreasing water potentials, (C) heat-shock treatments applied for 1 min and (D) storage for up to 24 months. Large coloured points show model-predicted germination percentages and black error bars indicate 95% confidence intervals; smaller transparent points represent individual replicates (n = 4). Different letters indicate significant differences among treatments based on Šidák-adjusted pairwise comparisons (P < 0.05). Germination percentages were calculated based on the number of viable seeds in each replicate. Seeds in the water potential, heat-shock and storage experiments were incubated for germination at 25 °C under a 12-h photoperiod.

**Table 1.** Viability of *Xyris paradisiaca seeds* following exposure to different temperatures, water potentials, heat shocks and storage durations. Values are mean viability percentages ± SE (n = 4). Different letters within each experiment indicate significant differences among treatments based on Šidák-adjusted pairwise comparisons following binomial GLMs.

| Experiment | Treatment | Seed viability (%) |
|---|---|---|
| **Temperature** | 10 °C | 78.8 ± 1.8 BCD |
|  | 15 °C | 99.0 ± 1.0 A |
|  | 20 °C | 92.7 ± 1.1 ABC |
|  | 25 °C | 97.0 ± 1.9 A |
|  | 30 °C | 94.1 ± 1.1 AB |
|  | 35 °C | 93.9 ± 2.6 AB |
|  | 40 °C | 74.4 ± 3.8 D |
|  | 45 °C | 77.0 ± 3.0 CD |
| **Water potential** | 0 MPa | 97.0 ± 1.9 A |
|  | −0.2 MPa | 91.2 ± 4.4 AB |
|  | −0.4 MPa | 96.0 ± 1.6 A |
|  | −0.6 MPa | 99.0 ± 1.0 A |
|  | −0.8 MPa | 99.0 ± 1.0 A |
|  | −1.0 MPa | 81.0 ± 2.5 B |
| **Heat shock** | Control | 97.0 ± 1.9 A |
|  | 80 °C | 92.0 ± 1.6 A |
|  | 100 °C | 92.0 ± 2.3 A |
|  | 120 °C | 92.0 ± 3.7 A |
|  | 150 °C | 87.0 ± 5.0 A |
|  | 200 °C | 97.3 ± 1.3 A |
| **Storage** | Fresh | 97.0 ± 1.9 A |
|  | 3 months | 82.0 ± 4.8 B |
|  | 6 months | 96.0 ± 2.8 AB |
|  | 9 months | 95.0 ± 2.5 AB |
|  | 12 months | 96.2 ± 1.6 A |
|  | 24 months | 96.4 ± 0.11 A |

Water potential significantly affected germination percentage (df = 5, □^2^ = 83.01, p < 0.001). Germination remained statistically similar to the control (98.5%) down to –0.6 MPa, but declined significantly to 78.1% at –0.8 MPa and to 40.9% at –1.0 MPa (Fig. 2B). Viability remained high (≥91.2%) and statistically similar to the control from −0.2 to −0.8 MPa, but declined to 81.0% at −1.0 MPa (Table 1). The best-supported hydro-time model indicated a median base water potential of Ψb = –1.03 MPa and a hydro-time constant of θ□ = 4.01 MPa d.

Heat-shock temperature had a significant overall effect on germination percentage (df = 5, □^2^ = 13.00, p = 0.023). However, no individual pairwise comparison was significant after Šidák adjustment (p_adj_ > 0.05 for all comparisons). Similarly, heat-shock temperature did not significantly affect seed viability (df = 5, □^2^ = 9.07, p = 0.106), which remained high across treatments (Table 1). Germination remained relatively high across treatments, ranging from 82.61% at 100 °C to 98.97% in the nonheated control (Fig. 2C).

*X. paradisiaca* seeds were desiccation-tolerant. Mean germination was 96.15% in fresh seeds and 100% after desiccation, resulting in a VLI = −0.04, which classified the seeds as desiccation-tolerant according to the adopted VLI criterion and indicated no detectable loss of germination capacity following drying. Storage time significantly affected viability-adjusted germination (df = 5, O^2^ = 32.57, p < 0.001). Germination declined from 98.5% in fresh seeds to 78.9% after 3 months of storage, but subsequently remained high, reaching 98.5%, 93.3%, 99.5% and 96.4% after 6, 9, 12 and 24 months, respectively. Germination after 3 months was significantly lower than in fresh and 6-month-stored seeds, whereas no other pairwise differences were detected after Šidák adjustment. Seed viability was also significantly affected by storage time (df = 5, □^2^ = 24.08, p < 0.001), declining from 97% in fresh seeds to 82% after 3 months but remaining ≥95% from 6 to 24 months. Thus, despite the lower values observed after a 3-mo storage period, neither germination nor viability showed a progressive decline over the 24-month storage period (Fig. 2D; Table 1).

## DISCUSSION

The regeneration niche represents a critical component of a species’ ecological niche because environmental conditions during germination and establishment can determine where successful recruitment occurs (Grubb, 1977). Accordingly, the breadth of the germination niche may contribute to species’ ecological breadth and geographical ranges, with narrow germination requirements potentially restricting the conditions under which recruitment can occur (Donohue *et al*., 2010). In X. paradisiaca, however, we found a striking contrast between its highly restricted geographical distribution and its broad physiological germination niche. Germination showed a strict light requirement but remained high across a wide thermal range, with thermal-time modelling predicting a germination window from approximately 9 to 46 °C and an optimum near 33 °C. Seeds also tolerated reduced water availability, maintaining high germination down to −0.6 MPa and exhibiting a median base water potential of approximately −1.03 MPa. Likewise, germination remained relatively high following brief heat exposure, with no individual heat-shock treatment differing significantly from the nonheated control. Seeds additionally showed favourable short-term storage behaviour under ambient laboratory conditions. Together, these results indicate that, although light availability may strongly constrain the microsites suitable for germination, the highly restricted distribution of X. paradisiaca cannot readily be attributed to narrow thermal or hydric germination requirements. Instead, its distribution is likely shaped by additional environmental filters, habitat requirements or demographic processes operating beyond the germination stage.

The absolute light requirement for germination observed in *X. paradisiaca* seeds agrees with previous results from studies with congeneric species (Garcia *et al*., 2020). Light-dependent germination is commonly associated with small-seeded species, such as *X. paradisiaca*, whose limited reserves restrict successful emergence from deeper soil layers. Light can therefore act as an environmental cue indicating that seeds are sufficiently close to the soil surface for successful seedling emergence and establishment (Milberg *et al*., 2000; Pearson *et al*., 2002; Aud and Ferraz, 2012). By preventing germination following burial, this response may also contribute to the formation of persistent soil seed banks, as reported for other *Xyris* species (Garcia *et al*., 2020). Whether *X. paradisiaca* forms a persistent soil seed bank, however, remains to be tested through field burial experiments.

Herbaceous species from the Cerrado typically germinate between 20 and 30 °C (Zaidan and Carreira, 2008), yet *Xyris* from *campos rupestres* generally germinate between 15 and 30 °C, with optimal germination temperatures between 20 and 30 °C (Garcia *et al*., 2020). However, *X. paradisiaca* maintained high germination percentages across a comparatively broad range of temperatures, from 15 to 40 °C, with thermal-time modelling predicting even larger thermal ranges. Seed viability, however, declined outside the 15–35 °C range, reaching only 74.4% at 40 °C. This indicates that, although 40 °C remained permissive for germination of viable seeds, prolonged exposure to high temperatures may constrain recruitment. Notably, the broad thermal niche of *X. paradisiaca* despite its restricted distribution agrees with previous evidence showing that geographical range size is not necessarily related to germination niche breadth (Thompson and Ceriani, 2003; Giorni *et al*., 2018; Ordóñez-Parra *et al*., 2025). Indeed, *X. paradisiaca* maintained high germination at 40 °C, whereas the observed germination niches of several *Xyris* species from *campos rupestres* in south-eastern Brazil extended at most to 35 °C (Garcia *et al*., 2020). At the lower end of its thermal niche, the estimated base temperature of *X. paradisiaca* was similar to the lowest value previously reported for the genus (T_b_ = 9.0 °C in *X. asperula*) (Oliveira *et al*., 2021). The restricted distribution of *X. paradisiaca* may therefore depend more strongly on environmental filters acting during pre- or post-germination stages than on thermal requirements for germination alone, as also suggested for other species from *campo rupestre* (Soares da Mota and Garcia, 2013). At the same time, the comparatively high upper thermal limit of X. paradisiaca may reflect the warmer thermal conditions of Central Brazil relative to south-eastern Brazil, although the decline in viability at 40 °C suggests that temperatures near the upper end of this range impose physiological costs even when germination remains high.

The effects of reduced water availability on germination remain comparatively understudied in Cerrado species, particularly among herbaceous taxa (Ordóñez-Parra *et al*., 2026). Recent studies, however, have shown that herbaceous species from the Cerrado tend to be more sensitive to reduced water potentials than shrubs (Cruz-Júnior *et al*., 2026). Still, among Cerrado grasses, wet-grassland species can maintain higher germination percentages at lower water potentials than those from dry grasslands (Souza *et al*., 2022). In contrast to the generally high sensitivity reported for Cerrado ground-layer species, *X. paradisiaca* maintained >40% germination at -1.0 MPa, whereas most ground-layer species studied by Cruz-Júnior *et al*., (2026) exhibited <10% germination at –0.9 MPa. This tolerance was also reflected in the hydro-time model, which estimated a median base water potential of Ψ_b_ = –1.03 MPa. This value falls within the range of soil water potentials experienced in *campo úmido* habitats, where values can decline to approximately –1.5 MPa during the dry season (Ramos *et al*., 2017). Thus, our model suggests that a substantial fraction of the seed population can germinate across much of the hydric variation experienced in its natural habitat, resulting in broad opportunities for recruitment. Together with its broad thermal tolerance, these findings provide further evidence that X. paradisiaca has a broad germination niche despite its highly restricted geographical distribution. Seed viability remained >90% from the control to −0.8 MPa, even though germination had already declined significantly at −0.8 MPa. These results indicate that moderate reductions in water potential primarily inhibited germination rather than compromising seed survival, as previously shown for other Cerrado ground-layer species (Cruz-Júnior et al., 2026). At −1.0 MPa, however, viability declined significantly to 81%, suggesting that prolonged exposure to more severe water limitation may begin to compromise embryo survival, even though most seeds remained viable.

The capacity to withstand limited water availability is further supported by results from our desiccation tolerance experiments that showed that seeds are desiccation tolerant and are capable of germinating after drying. Altogether, these responses suggest that *X. paradisiaca* seeds tolerate both low external water potentials and substantial dehydration with comparatively limited loss of viability. Such tolerance may allow seeds to remain viable during periods of soil water deficit and retain the potential to germinate upon the return of precipitation at the onset of the rainy season. Studies of *Xyris* species from *campos rupestres* show that imbibed seeds can retain high germinability after subsequent drying, suggesting that seeds may withstand interrupted hydration caused by isolated rainfall events during the dry season or by dry spells during the rainy season (Oliveira *et al*., 2018). However, the capacity of seeds to resume germination following transient water limitation or drying after imbibition remains poorly characterised among Cerrado species, warranting further investigation in *X. paradisiaca* and other ground-layer taxa.

Seeds of our study species were not only tolerant of temperature and moisture gradients but also of heat shock, with seeds subjected to a 1-min oven treatment at 200 °C having germination and viability similar to those of nonheated seeds. Studies with congeneric species indicate a high tolerance to heat shock, with germination remaining unaffected at temperatures up to 150 °C (Oliveira *et al*., 2018). Notably, although soil-surface temperatures during Cerrado fires can exceed 400 °C, heating decreases sharply with depth, with temperatures measured 1 cm below the soil surface ranging from approximately 29 to 131 °C during experimental fires (Zupo *et al*., 2022). This heat tolerance is consistent with a widespread post-fire regeneration strategy among Cerrado ground-layer species, in which vegetative resprouting co-occurs with the production of heat-tolerant propagules, a combination that has also been documented for *Xyris* (Pilon *et al*., 2021; Zupo *et al*., 2021; Daibes *et al*., 2022). Beyond heat tolerance, fire may also promote recruitment through chemical cues. In *X. paradisiaca*, smoke-water exposure increased germination percentage and reduced mean germination time (Motta *et al*., 2024), while studies of other Xyridaceae and campo rupestre species have shown that smoke can accelerate and synchronise germination (Le Stradic *et al*., 2015; Fernandes *et al*., 2021). These responses indicate that smoke-derived compounds can act as post-fire germination cues for *Xyris* species. Together, these findings suggest that fire may influence the regeneration of *X. paradisiaca* through complementary processes. That is, seeds can survive the short-duration heating associated with fire, while subsequent exposure to smoke-derived compounds may promote and accelerate germination in the post-fire environment.

*X. paradisiaca* seeds showed no progressive loss of germinability or viability over 2 years of storage under ambient laboratory conditions, indicating favourable short- to medium-term storage potential. Although both germination and viability were lower after 3 months, values were again high from 6 to 24 months, suggesting that the decline observed at 3 months did not represent a sustained deterioration with increasing storage duration, but rather variation associated with the 3-month seed lot. Importantly, this performance under simple storage conditions suggests that seeds may be temporarily stored using relatively low-technology methods, providing a readily applicable option for local seed collectors, conservationists and restoration practitioners. Such flexibility could facilitate the accumulation, handling and distribution of seed lots for propagation and restoration activities, particularly where access to refrigerated storage is limited. The maintenance of high germinability after 24 months also indicates potential for longer-term ex situ conservation (De Vitis *et al*., 2020). This potential is further supported by the desiccation tolerance observed in our experiment and by storage-behaviour records for the genus, with seven *Xyris* species currently reported as orthodox in the Seed Information Database (https://ser-sid.org/; last access: August 27, 2026). The small size and low mass and moisture content characteristic of *Xyris* seeds may also favour storage under dry conditions (Garcia *et al*., 2020). Nevertheless, desiccation tolerance and 2 years of survival under ambient conditions are insufficient to establish orthodox storage behaviour in *X. paradisiaca*. Further assessments of seed longevity across a range of controlled moisture contents and storage temperatures are essential to determine appropriate protocols for long-term ex situ seed banking (De Vitis *et al*., 2020).

Overall, our results indicate that *X. paradisiaca* combines broad abiotic germination tolerances with desiccation tolerance and favourable storage behaviour despite its highly restricted geographical distribution. These traits indicate considerable potential for seed-based conservation, ecological restoration and nursery propagation. This potential is particularly relevant in the PNCV region, where community-based native-seed networks have become an important source of propagules for restoration while generating income for local communities (Schmidt *et al*., 2019). The capacity of *X. paradisiaca* seeds to retain high germinability under simple ambient storage conditions may facilitate their incorporation into these seed-supply chains by allowing collectors to maintain seed lots using relatively simple storage methods before distribution or use. Incorporating *X. paradisiaca* into such initiatives, where legally and ecologically appropriate, could also contribute to diversifying restoration seed mixtures, as forbs represent a major component of Cerrado plant diversity but remain disproportionately underrepresented in the native-seed market (Silva *et al*., 2022). Because germination requires light, however, sowing methods should avoid deep seed burial, as 5 cm burial was enough to completely inhibit germination in other *Xyris* species (Oliveira *et al*., 2017; Oliveira and Garcia, 2019). Ensuring adequate light exposure during sowing, together with the broad thermal and hydric germination tolerances identified here, may therefore facilitate propagation under relatively simple nursery conditions. Indeed, successful propagation under common, low-cost nursery conditions has been demonstrated for numerous endemic species from Brazilian open ecosystems (Faria *et al*., 2025).

Developing propagation protocols may have benefits beyond restoration. Harvesting of flowering individuals for the everlasting-flower trade is among the threats reported for *X. paradisiaca* (Martinelli and Moraes, 2013), yet everlasting-flower harvesting forms part of a culturally and economically important Cerrado value chain involving Xyridaceae and other native plants (Neves *et al*., 2026). Cultivation could therefore provide an alternative source of plants for ornamental use while reducing dependence on harvesting threatened wild populations. Moreover, the ornamental potential of endemic plants from Brazilian open ecosystems remains largely untapped, making cultivation of *X. paradisiaca* a potential avenue for conservation gardening and complementary *ex situ* conservation (Faria *et al*., 2026). More broadly, our findings show that the highly restricted distribution of this Endangered endemic is not driven by a narrow physiological germination niche, while identifying seed traits that can inform its conservation, propagation and restoration.

## ACKNOWLEDGMENTS

We thank the Instituto Chico Mendes de Conservação da Biodiversidade (ICMBio) for providing the legal permits for conducting this research, and Eduardo D. Lozano for confirming the species identification. We also thank Fernando A. O. Silveira for his comments on a previous version of the manuscript. ChatGPT (GPT-6) was used for language editing of the text written by the authors, including grammar, clarity and readability; not to generate original scientific content, analyses or interpretations. The authors reviewed all edited text and remain fully responsible for the content of the manuscript.

## AUTHOR CONTRIBUTIONS

CAO-P conceived the study. CAO-P, MB, LA-G, RF, TEBA and CAC collected the data, which were analysed by CAO-P. All authors contributed to the writing process, proofread and corrected the manuscript.

## DATA ACCESSIBILITY STATEMENT

All the data and R code used in this study will be available on a GitHub repository (https://github.com/caordonezparra/xyris_paradisiaca_germination) upon publication.

